# Comprehensive mass spectrometry-guided plant specialized metabolite phenotyping reveals metabolic diversity in the cosmopolitan plant family Rhamnaceae

**DOI:** 10.1101/463620

**Authors:** Kyo Bin Kang, Madeleine Ernst, Justin J. J. van der Hooft, Ricardo R. da Silva, Junha Park, Marnix H. Medema, Sang Hyun Sung, Pieter C. Dorrestein

## Abstract

Plants produce a myriad of specialized metabolites to overcome their sessile habit and combat biotic as well as abiotic stresses. Evolution has shaped specialized metabolite diversity, which drives many other aspects of plant biodiversity. However, until recently, large-scale studies investigating specialized metabolite diversity in an evolutionary context have been limited by the impossibility to identify chemical structures of hundreds to thousands of compounds in a time-feasible manner. Here, we introduce a workflow for large-scale, semi-automated annotation of specialized metabolites, and apply it for over 1000 metabolites of the cosmopolitan plant family Rhamnaceae. We enhance the putative annotation coverage dramatically, from 2.5 % based on spectral library matches alone to 42.6 % of total MS/MS molecular features extending annotations from well-known plant compound classes into the dark plant metabolomics matter. To gain insights in substructural diversity within the plant family, we also extract patterns of co-occurring fragments and neutral losses, so-called Mass2Motifs, from the dataset; for example, only the Ziziphoid clade developed the triterpenoid biosynthetic pathway, whereas the Rhamnoid clade predominantly developed diversity in flavonoid glycosides, including 7-*O*-methyltransferase activity. Our workflow provides the foundations towards the automated, high-throughput chemical identification of massive metabolite spaces, and we expect it to revolutionize our understanding of plant chemoevolutionary mechanisms.

## INTRODUCTION

Specialized metabolites, also called secondary metabolites or natural products, are molecules produced by all higher plants; and deployed for the survival in a competitive environment (Hartmann, 2007). The chemical diversity in the plant kingdom has been accumulated over evolutionary time. Therefore, the distribution of specialized metabolites across the plant kingdom is an important aspect of phenotyping that can, for example, provide us with insights about the evolution of biosynthetic pathways (Wink, 2003). However, directly assessing plant chemical diversity is extremely challenging and several bottlenecks have limited large-scale studies investigating the evolutionary history of plant specialized metabolism. Chemotaxonomic studies assessing the relationship between plant morphological characters and chemical composition have largely depended on literature surveys, which do not only require a large investment in time and labor but also involve a lot of biases. For example, there is a general emphasis towards single isolated plant specialized metabolites that exhibit biological activities with pharmaceutical interest (Harvey, 2008); also, it is common practice not to publish chemical structural information of molecules, which do not exhibit structural novelty or biological activities of interest. Experimentally assessing chemical diversity among plants has been limited by the inability to automate chemical structural characterization, something that is still an inherently slow and largely manual process that needs expert knowledge.

Here, we introduce a scalable workflow to digitize diversity and distribution of plant specialized metabolites using mass spectrometry (MS) in combination with a series of computational mass spectrometry data analysis tools. In theory, tandem mass spectrometry (MS/MS) contains a lot of information that can be used to gain structural insight into the molecules that are detected (Ernst *et al.*, 2014). However, annotation, classification and identification of metabolites that are detected by MS is still a significant obstacle in plant metabolomics workflows, in contrast to high-throughput characterization of DNA, RNA, and proteins where annotation and classification have become much more routine even when MS/MS is employed (Nakabayashi and Saito, 2013). Computational tools such as *in silico* fragmentation predictors and combinatorial fragmentators (Allen *et al.*, 2014, Duhrkop *et al.*, 2015, Ruttkies *et al.*, 2016, da Silva *et al.*, 2018) and molecular networking (Watrous *et al.*, 2012, Wang *et al.*, 2016) combined with library matching to reference spectra have enabled automated chemical structure annotations in recent years. Even with those advances, only ~2–5 % of the MS/MS spectra can be annotated in an experiment (da Silva *et al.*, 2015, Wang *et al.*, 2016, Aksenov *et al.*, 2017). To enhance the coverage of putative annotation on MS/MS spectra, we developed a scalable semi-automated approach towards the characterization of plant specialized metabolites by integrating several computational MS/MS data analysis methods (Figure 1). Most previous *in silico* annotation methods focused on putative identification of individual molecules of interest; in contrast, our workflow putatively annotates molecular families (groups of molecules having common chemical scaffolds; Nguyen et al., 2013). Combining information on both full structures and predicted substructures of multiple molecules, and the motifs and fragmentation patterns associated with these, allows our workflow to greatly extend the number of spectra that can be annotated. We developed a scalable semi-automated approach towards the chemotaxonomic characterization of plants, and demonstrate the efficiency of our workflow on a unique collection of extracts of 70 species from the Rhamnaceae family. Rhamnaceae is a cosmopolitan plant family of ~50 genera and 900 species (Richardson et al., 2004). Rhamnaceae species are known for their exceptional morphological diversity and high genetic variation, likely as evolutionary consequences associated with its wide geographic distribution and many different habitats (Hardig *et al.*, 2000, Hauenschild *et al.*, 2016a, Hauenschild *et al.*, 2016b). Although there are some family-specific metabolites such as ceanothane-type triterpenoids (Kang *et al.*, 2016) and cyclopeptide alkaloids (Tuenter *et al.*, 2017), little chemistry is known from this family. We employed next generation metabolomics data analysis strategies to provide structural insight into hundreds of specialized metabolites both at the level of chemical class and diversified scaffolds.

**Figure 1.**
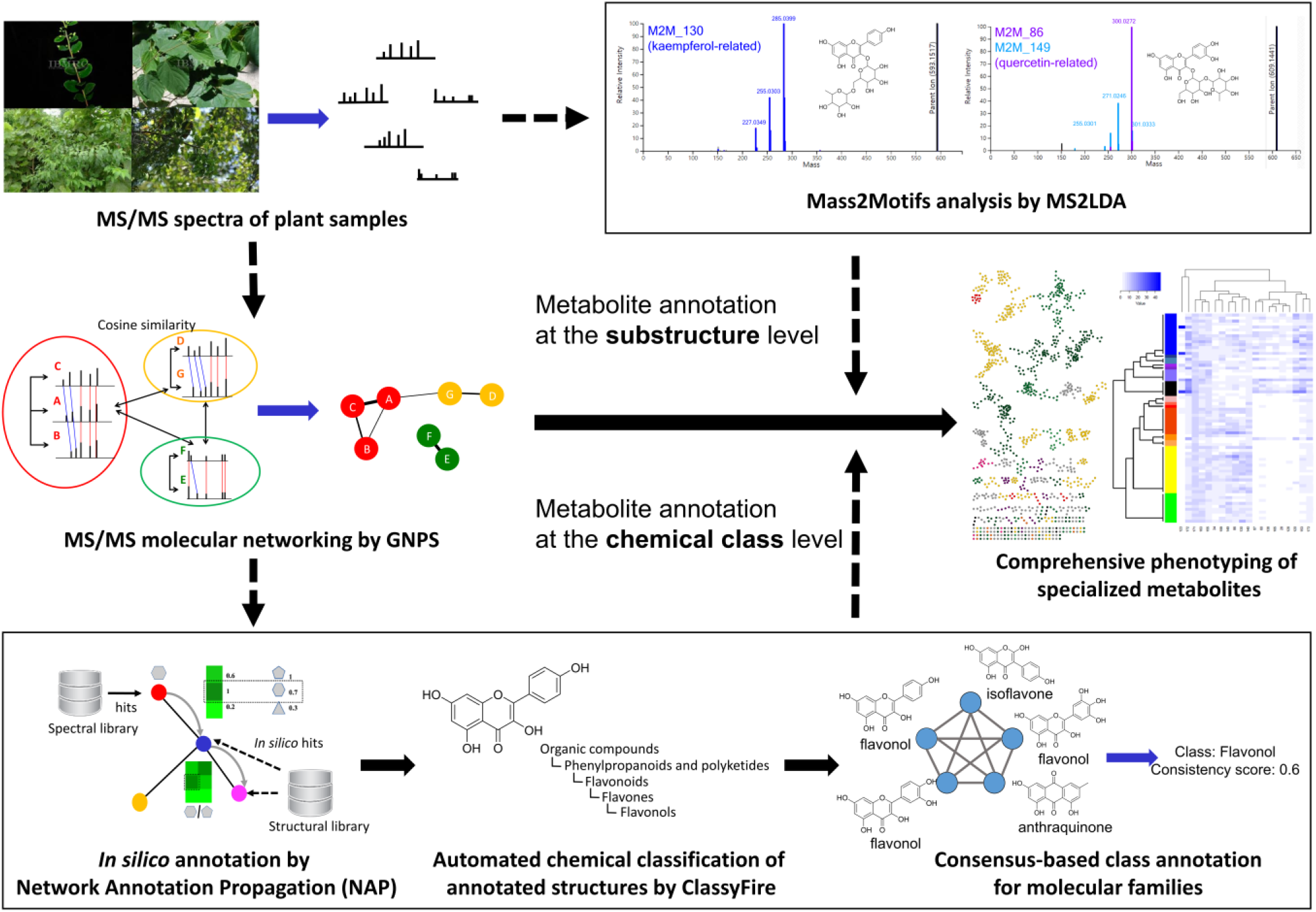
Schematic data analysis workflow for comprehensive plant specialized metabolites phenotyping using MS/MS. MS/MS spectra are analyzed for spectral similarity and visualized as a molecular network, which clusters similar spectra as “molecular families”. Network annotation propagation (NAP) provides *in silico* annotation candidates for individual spectra. These candidates are chemically classified using ClassyFire, then molecular families are putatively annotated based on the most predominant chemical classes per molecular family. Meanwhile, the distribution of co-occurring fragments and neutral losses (Mass2Motifs) are analyzed by MS2LDA, and these provide information about substructure diversity and distribution between samples.

## RESULTS AND DISCUSSION

### The Rhamnaceae chemical space

To take an inventory of the Rhamnaceae plant family, we submitted LC–MS/MS data from 70 representative Rhamnaceae species extracts to mass spectral molecular networking through the Global Natural Products Social Molecular Networking (GNPS) web platform (https://gnps.ucsd.edu)(Wang *et al.*, 2016). The resulting molecular network consisted of 2,268 mass spectral nodes organized into 141 independent molecular families (two or more connected nodes of a graph; Nguyen *et al.*, 2013). We investigated chemical diversity in relation to the most recent phylogenetic study (Sun *et al.*, 2016). Based on this phylogenetic hypothesis, our 70 Rhamnaceae species spanned 15 genera. These 15 genera are further grouped into two major phylogenetic clades, the Rhamnoid clade, comprising 8 genera and the Ziziphoid clade, comprising a total of 7 genera (Richardson *et al.*, 2000, Sun *et al.*, 2016) (Figure 2(b)). Phylogenetically closely related genera were assigned similar colors, so that phylogenetic relationships could be visualized on the mass spectral molecular network (Figure 2(a)). We observed that specialized metabolite classes tend to be constrained to specific taxa. For example, more than 90% of the metabolites within the molecular families **A** and **B** were predominantly found in one phylogenetic clade; Rhamnoid for **A** and Ziziphoid for **B** (Figure 2(c)). Furthermore, molecular family **C** exhibits molecules found in representatives of both clades and several genera, suggesting widespread occurrence of certain metabolite classes within Rhamnaceae species. We further detected species or genera-specific chemical analogues. For example, some spectral nodes within the molecular family **B** are unique to the genus *Gouania*, while the others are found only in *Colubrina* species. This finding reveals the presence of closely related yet different chemical structures across members of these two genera.

**Figure 2.**
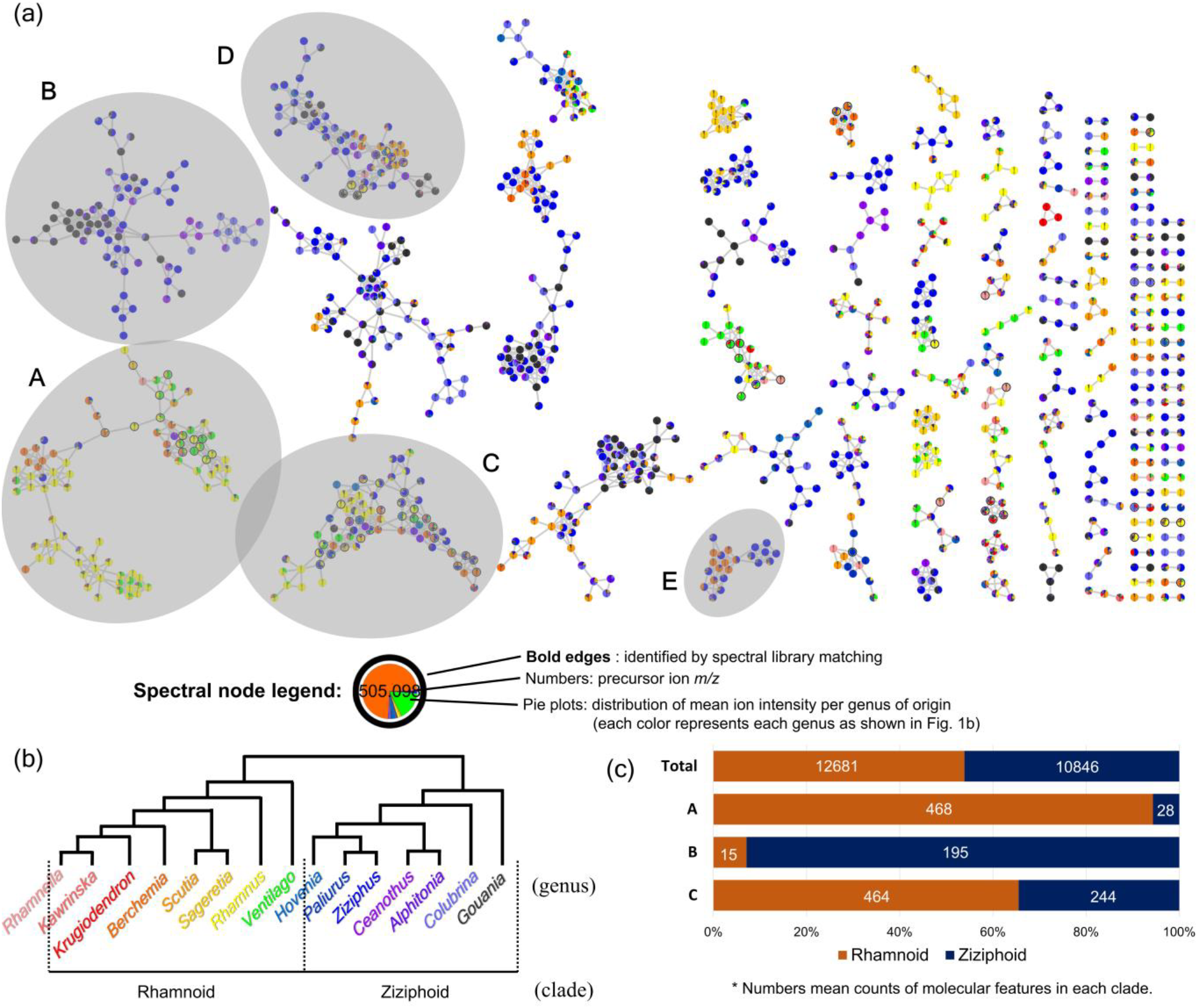
The Rhamnaceae molecular network and mass spectrometry detected chemical space. (a) Global Rhamnaceae mass spectral molecular network with nodes colored according to the mean ion intensity per genus of origin. Molecular families **A**–**E** (**A**, various phenolics; **B**, triterpene glycosides; **C**, flavone O-glycosides; **D**, triterpene esters; **E**, cyclopeptide alkaloids) are highlighted. (b) Schematic representation of Rhamnaceae phylogenetic tree retrieved from Richardson *et al.*, 2000 and Sun *et al.*, 2016. Phylogenetically closely related genera were assigned similar colors. (c) Distribution of metabolites within the Ziziphoid and Rhamnoid clades across the global mass spectral molecular network and molecular families **A**, **B**, and **C.** Differential abundance was assessed based on binary counts of MS1 ions in each species.

### Metabolite annotation at the subclass level

The MS/MS spectral library search through GNPS as described in the Experimental section resulted in 51 hits to reference MS/MS spectra. These are level 2 or 3 annotations according to the 2007 metabolomics standards initiative (MSI) (Sumner *et al.*, 2007). This is about 2.5 % of the observed Rhamnaceae chemical space. Most of the library hits belong to the molecular families of flavonoid glycosides (e.g. **A** and **C**), because experimental MS/MS spectra in public spectral libraries are not equally distributed across different chemical classes. There is a strong bias in the public libraries towards commercially available molecules and more abundant metabolites as this facilitates isolation and structure elucidation. To amplify the chemical knowledge that we can obtain from the data, we applied *in silico* structure prediction (network annotation propagation, NAP) to obtain *in silico* fragmentation-based metabolite annotation candidates from relevant compound databases, through reranking candidate molecular annotations based on the network topology (da Silva *et al.*, 2018). Except for 87 MS/MS spectra, NAP assigned candidate structures to the majority of the nodes. Matching failures are usually a result of lack of candidate structures within the corresponding compound libraries.

Molecular networking utilizes spectral similarity to group metabolites with the implicit assumption that similar molecular structures will generate similar fragmentation spectra; thus, molecular families comprising structurally similar molecules can likely be interpreted as distinct chemical classes. Based on this hypothesis, structures annotated by NAP were classified based on their chemical scaffolds using ClassyFire (Djoumbou Feunang *et al.*, 2016). ClassyFire assigned chemical structures to a chemical ontology consisting of up to 11 different levels, and the most frequent consensus classifications per molecular family were retrieved (Figure 3(a)). Reliability of the ClassyFire analysis was validated using two different scores. At first, the ratio of nodes returning any database hit from NAP, the coverage score, was calculated for all molecular families. 90.78 % of all molecular families within our global network showed a coverage score of over 0.7, indicating high structural library coverage of our samples (Figure S10). Meanwhile, the consistency score, defined as the % of nodes that make up a molecular family, indicates how coherent the ClassyFire classifications are (Figure 3(a)). The NAP annotated molecular families varied in their consistency. NAP is dependent on structural library hits, and many molecules that can be detected are not covered in structural libraries. Some network clusters consist of different classes of metabolites, while others show higher coherence for their identified structures. For example, the ClassyFire result revealed that molecular families **A** and **C** were primarily composed of flavonoid glycosides. The consistency score of **A** was 0.394, indicating that 39.4% of all structural matches in **A** were classified as flavonoid glycosides; on the other hand, molecular family **C** showed a score of 0.688. Manual inspection of **A** and **C** support the classification results as diverse subgroups of related phenolic (e.g. flavonoids, anthraquinones, and naphthopyrones) glycosides were found in **A**, while most nodes in **C** were annotated as flavonoid glycosides (Data S1, Supporting Information). This indicates that the annotation of chemical classes could be a broad strategy for exploring the chemical space and diversity of large metabolomics datasets.

**Figure 3.**
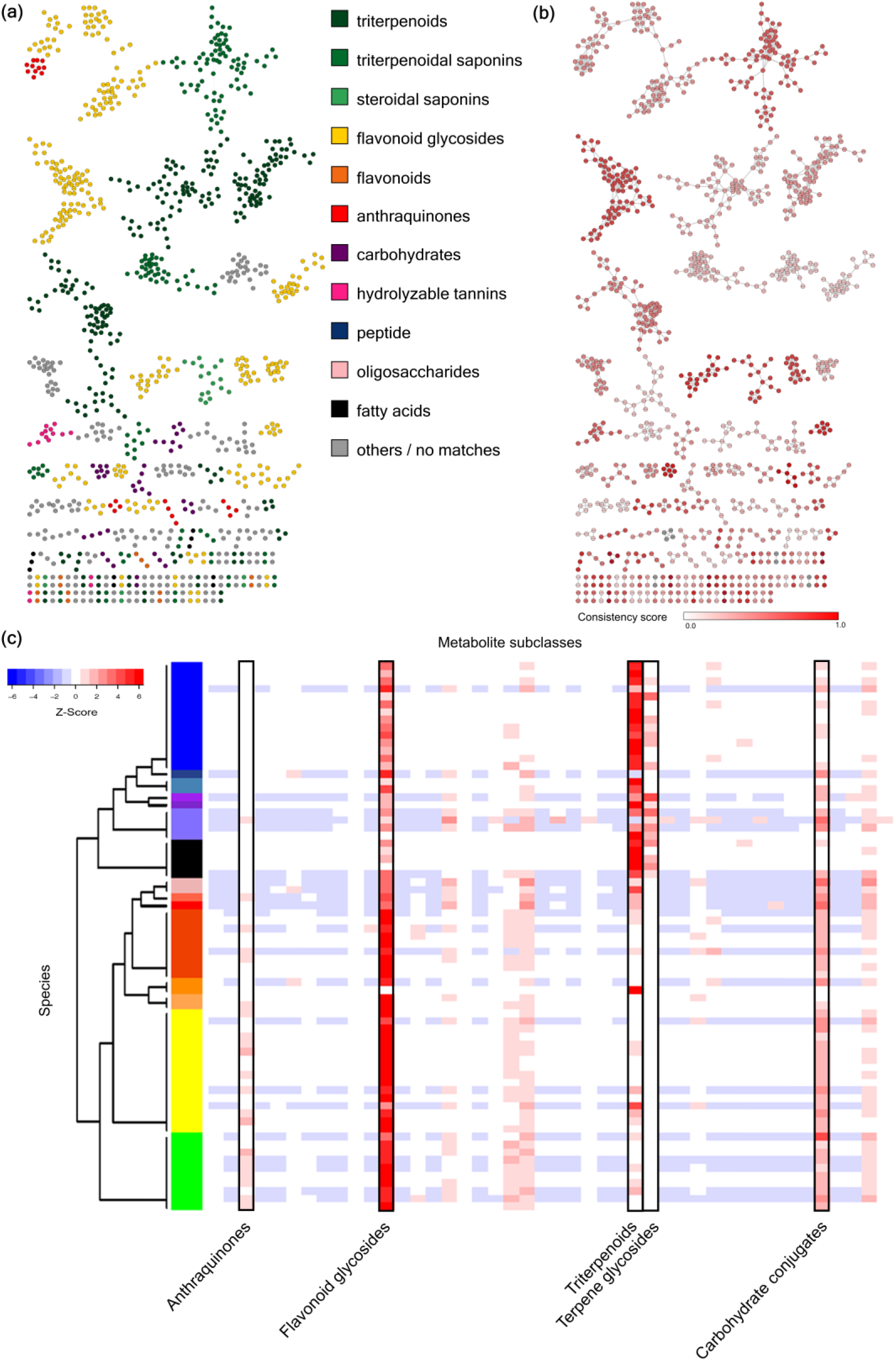
Structural annotation of Rhamnaceae specialized metabolites at the chemical subclass level. (a) Chemical structures annotated by NAP were automatically classified for their chemical scaffolds using ClassyFire, and the most frequent consensus classifications per network cluster were retrieved to assign putative chemical subclass annotation to each molecular family. (b) The ClassyFire consistency score which indicates the coherence of the ClassyFire chemical classification across each molecular family was calculated to estimate the accuracy of putative annotations. (c) Heatmap of the normalized putatively identified molecular features illustrating distribution of specialized metabolite classes across 70 Rhamnaceae species. Each column represents a specialized metabolite class while each row represents a species. For visualization purposes, a few differentially expressed chemical classes are highlighted. The complete heatmap can be found in Figure S11.

Based on the putative chemical classification of molecular families, the normalized distribution pattern of different classes of metabolites were visualized as a heatmap (Figure 3(b)). On the y-axis, we plotted the samples, and on the x-axis the putative chemical classes. The colour scheme in the heatmap represents Z-scores per sample. It was revealed that Ziziphoid species exhibit various triterpenoids and triterpenoid glycosides, while Rhamnoid species show more diversified flavonoids, carbohydrates, and anthraquinones. However, most of chemical classes did not show very conserved patterns in specific genera or tribes, being suggestive of convergent evolution in specialized metabolism. This finding would corroborate with the extraordinary convergent genetic diversity of Rhamnaceae caused by their worldwide distribution, especially in Mediterranean-type ecosystems (Onstein *et al.*, 2015, Onstein and Linder, 2016).

### Metabolite annotation at the scaffold diversity level

Plant specialized metabolite profiles often show a pattern in which a few major metabolites occur widely in certain level of taxa, and those major compounds are accompanied by several minor derivatives (Wink, 2003). Although more than 200,000 natural products are known to be synthesized by plants, all of those are based on only a few biosynthetic pathways and key primary metabolites. Therefore, a small portion of metabolites tend to be observed universally across the plant kingdom, while minor derivatives of them show more specific distributions caused by independently evolved downstream pathways. Substructure recognition topic modeling (MS2LDA) (van der Hooft *et al.*, 2016) was applied on our MS/MS dataset for extraction of information on substructural diversity within each metabolite class. MS2LDA reveals patterns of co-occurring fragments and neutral losses (called Mass2Motifs) from multiple MS/MS spectra (van der Hooft *et al.*, 2016). 200 motifs were retrieved from the dataset with MS2LDA - of which we could annotate 25 with chemical substructures using the MS2LDAviz web app (Wandy *et al.*, 2017). Figure 4 visualizes the distribution of MS/MS spectra containing each Mass2Motif, which represents substructural diversity among the tested species. This provides insights about how scaffold diversity has evolved in this family. For example, Mass2Motif 179 which is related to rhamnetin (7-*O*-methylquercetin) is only observed in Rhamnoid species, while quercetin-related metabolites are observed across the entire family. It suggests that quercetin 7-*O*-methyltransferase is active only in Rhamnoid species, while it is silent or has not evolved in Ziziphoid species. Although we cannot validate this hypothesis due to low coverage of the Rhamnaceae genome (Liu *et al.*, 2014), our approach provides a very straightforward way to phenotype-based hypotheses within plant specialized metabolism.

**Figure 4.**
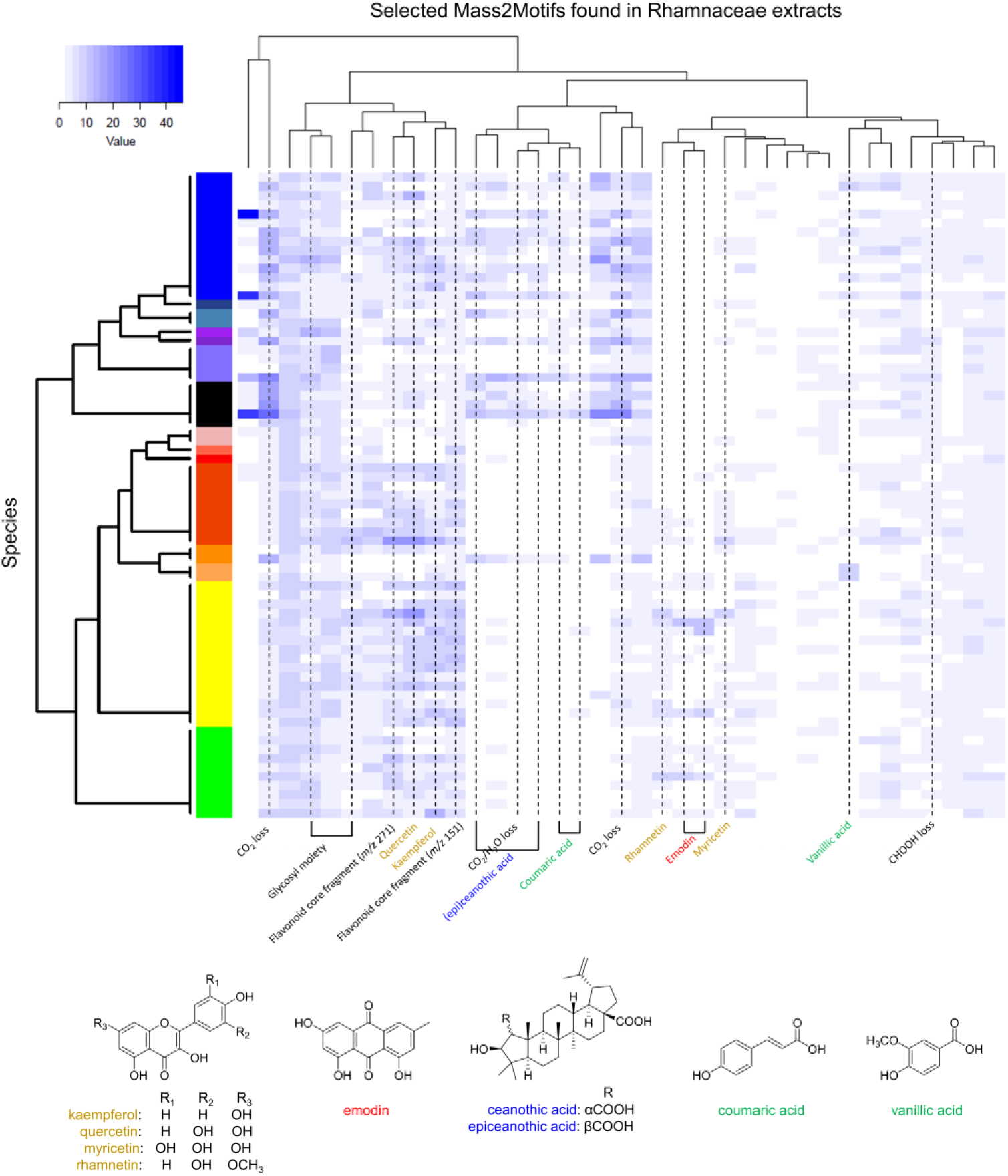
Heatmap of the molecular features illustrating distribution of different Mass2Motifs across 70 Rhamnaceae species (The counts of molecular features related to Mass2Motifs were filtered with the probability > 0.3). Each column represents a Mass2Motif while each row represents a species. Selected substructures related to annotated Mass2Motifs are highlighted drawn below the heatmap. The complete heatmap can be found in Figure S12.

Our workflow provides insights on plant specialized metabolism on a systemic level; however, both NAP and MS2LDA work on each individual spectrum. Thus, this workflow can also be exploited for the annotation of specific molecules of interests, which especially agrees with interests of natural product chemists. Figure 5(a) describes a summary of the metabolite annotations in molecular family **A**. It shows the synergism of using both NAP and MS2LDA for annotation of MS/MS spectra. Mass2Motif 164 could be annotated as rhamnocitrin (7-*O*-methylkaempferol)-related motif based on the putative annotation of node **1** as rhamnocitrin-3-*O*-rhamninoside, while rhamnetin-related Mass2Motif 179 were extracted from spectral nodes **2** (rhamnazin-3-*O*-rhamninoside), **3** (rhamnetin-3-*O-*rhamninoside), and **4** (rhamnetin-3- *O-*rutinoside). Distribution mapping of Mass2Motifs 40, 64,141, and 168 also revealed scaffold differences of emodin, norrubrofusarin, and torachrysone in MS/MS spectral nodes clustered as the molecular family **A** (Figure 5(a)). Figure 5(b) shows another example; molecular family **D** was putatively identified as a family of triterpene esters. Different phenolic moieties such as protocatchuate, vanillate, and coumarate were easily recognized in **D**, by analyzing the distribution of Mass2Motifs 28, 117, 120, and 191. We validated 8 molecular annotations classified as flavonoids, anthraquinones, triterpenoids, and peptides using reference standards, and all of them were confirmed as the correct structural annotation (Result S2, Supporting Information) thus promoting them to MSI level 1 identifications (spectra are available in GNPS public library – see Result S2). Therefore, we suggest that the workflow introduced in this article will enhance both the efficiency of dereplication, the process of identifying “unknown knowns” from complex mixtures, and illumination of the “unknown unknown” dark metabolic matter, both critical steps for the natural product drug discovery process.

**Figure 5.**
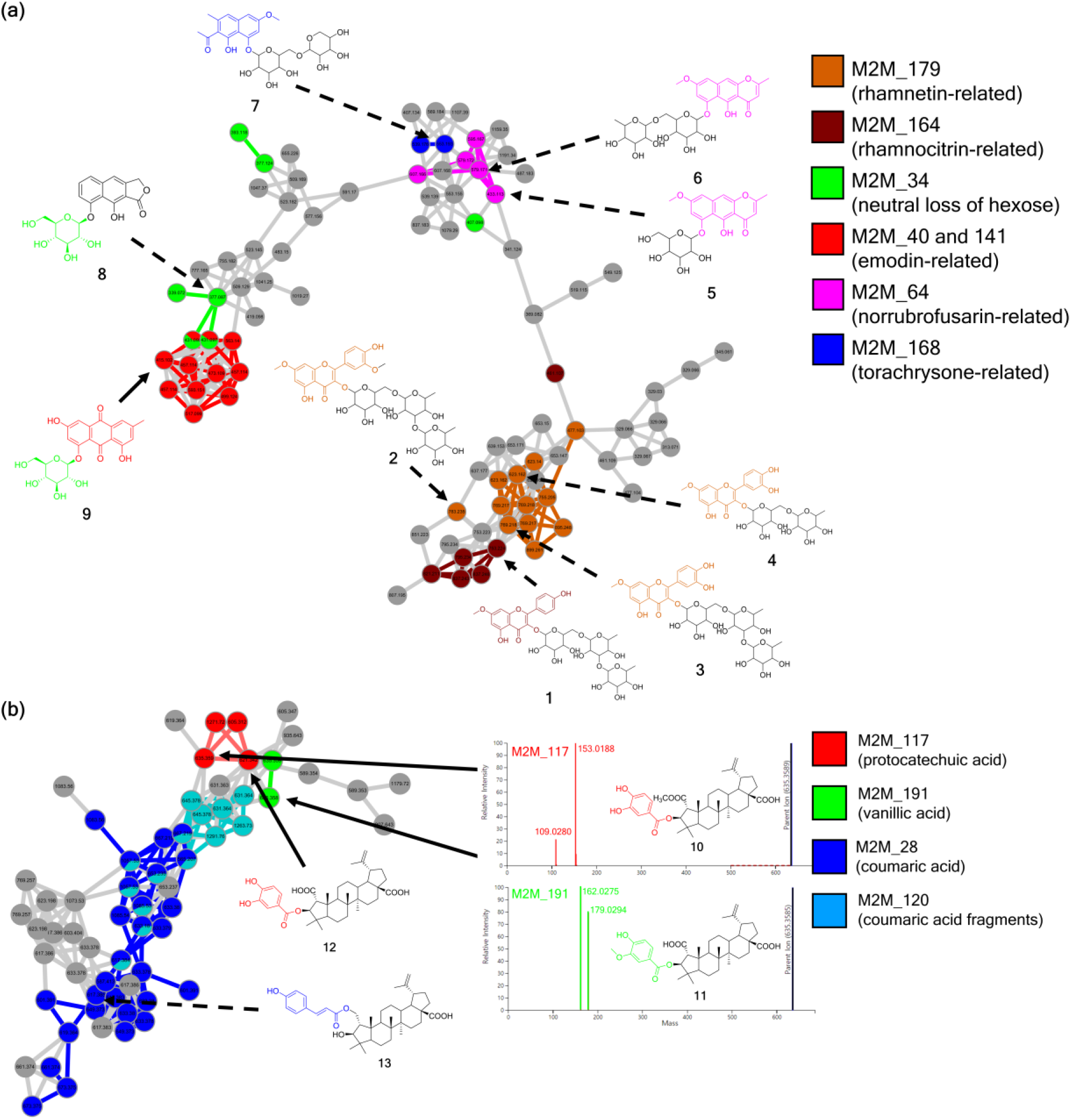
NAP/MS2LDA-driven metabolite annotation of (a) diverse phenolics (molecular family **A**) and (b) triterpene esters (molecular family **D**). Mass2Motifs, mapped with different colors of spectral nodes, reveal diversity of chemical scaffolds (a) and substructural moieties (b). Chemical structures drawn here are top-candidates suggested by the NAP analysis. Colored parts for structures represent the substructures related to Mass2Motifs. The annotated structures of compounds **9**–**12** were authenticated using reference standards.

## CONCLUSIONS

Although metabolomics is a rapidly growing discipline in plant science, its application is still relatively limited, compared to genomics or transcriptomics. The lack of a high-throughput annotation method is one of the major reasons for it. Using the integrative workflow based on MS/MS molecular networking, we were able to putatively annotate and classify metabolites in high-throughput. Although most annotations still need to be inspected and validated manually, we can reach consensus and higher confidence in chemical structural data interpretation by using two different, complementary computational approaches, MS2LDA and automated chemical classification of *in silico* annotated structures within the mass spectral molecular networks. Therefore, we expect that this workflow, in addition to expansion of public spectral databases coverages, *in silico* annotation tools’ accuracy improvement and comprehensive substructure annotation to accelerate the application of metabolomics approaches to plant biology. These advances are likely to reproduce what the GenBank (Benson *et al.*, 2018), Basic Local Alignment Search Tool (BLAST) (Altschul *et al.*, 1990) and Gene ontology broad usage (Ashburner *et al.*, 2000) did for genomics studies. Based on the putative annotations, we were able to characterize, analyze, and visualize the chemical space of the Rhamnaceae family, allowing us to digitize the diversity and distribution of metabolites. Considering that most species used in this study have not been engaged in any phytochemical studies, we expect that our method will accelerate chemical identification of uncharted plant metabolite space. There have been other approaches for accelerating plant metabolite identification, such as candidate substrate-product pair (CSPP) network (Morreel *et al.*, 2014), ISDB-molecular networking (Allard *et al.*, 2016), MatchWeiz (Shahaf *et al.*, 2016), or PlantMAT (Qiu *et al.*, 2016). However, all these approaches not only rely on compound database content like our approach but also previous knowledge such as reported phytochemical composition or metabolic pathways. Unfortunately, this information is hard to obtain for many taxa; as there still is a large number of plants whose metabolomic composition has never been investigated. Recent studies revealed that convergent evolution can lead to identical specialized metabolites biosynthesized through different unrelated pathways (Huang *et al.*, 2016, Zhao *et al.*, 2016). Therefore, researchers should consider the risk of drawing incorrect conclusions when they apply specialized metabolic pathways established in different plants. In this context, our approach has an advantage in digitizing and visualizing the chemical space of both previously investigated as well as uninvestigated plants, because it does not have any taxonomic bias using only MS/MS data and available molecular structures from compound databases. Hence, our approach facilitates metabolomics studies with massive datasets from uninvestigated species for botany, ecology, evolutionary biology, and natural products discovery. Currently the workflow is available for all users by using the scripts (available at https://github.com/DorresteinLaboratory/supplementary-Rhamnaceae), and the work is ongoing to wrap up the analysis workflow in one package minimizing the number of scripts needed to get to an enhanced molecular network.

## EXPERIMENTAL PROCEDURES

### Plant materials

Aerial parts of 70 Rhamnaceae plant species were collected in Cambodia, China, Costa Rica, Ecuador, Indonesia, Laos, Mongolia, Nepal, and Vietnam. Samples were extracted with MeOH or 95 % EtOH, after drying and pulverizing. Extraction solvents were immediately removed by freeze-drying, and the dried extracts were stored at − 20 °C until analyses. The samples were authenticated by collectors, and voucher specimens are deposited in the International Biological Material Research Center (IBMRC) of Korea Research Institute of Bioscience and Biotechnology (KRIBB), together with the extract library. Detailed location and date for collection are listed in Table S1.

### Liquid chromatography coupled tandem mass spectrometry (LC–MS/MS)

Dried extracts were re-dissolved in MeOH at a concentration of 5mg/mL and analyzed using an ACQUITY ultra-high performance liquid chromatography (UPLC) system (Waters Co., Milford, MA, USA) coupled to a Xevo G2 QTOF mass spectrometer (Waters MS Technologies, Manchester, UK) equipped with an electrospray ionization (ESI) interface. Chromatographic separation was performed on an ACQUITY UPLC BEH C_18_ (100 mm × 2.1 mm, 1.7 μm, Waters Co.) column eluted with a linear gradient of 0.1% formic acid in H_2_O (A) and acetonitrile (B) with increasing polarity (0.0 to 14.0 min, 10% to 90% B). The column was maintained at 40 °C, the flow rate was 0.3 mL/min, and the linear gradient elution was followed by a 3 min washout phase at 100% B and a 3 min re-equilibration phase at 10% B. Analyses of the extract samples (1.0 μL injected into the partial loop in the needle overfill mode) were performed in negative ion automated data-dependent acquisition (DDA) mode, in which full MS scans from *m/z* 100–1500 Da are acquired as MS1 survey scan (scan time: 150 ms) and then MS/MS scans for the three most intense ion follow (scan time: 100 ms). MS/MS acquisition was set to be activated when TIC of MS1 survey scan rose and switched back to survey scan after two scans of MS/MS. The ESI conditions were set as follows: capillary voltage 2.5 kV, con voltage 20 V, source temperature 120 °C, desolvation temperature 350 °C, cone gas flow 50 L/h, and desolvation gas flow 800 L/h. High-purity nitrogen was used as the nebulizer and auxiliary gas, and argon was used as the collision gas. Data were acquired in centroid mode, and the [M − H]^−^ ion of leucine enkephalin at m/z 554.2615 was used as the lock mass to ensure mass accuracy and reproducibility. Collision energy gradient was automatically set according to m/z values of precursor ions: 20 to 40 V for 100 Da to 60 to 80 V for 1500 Da.

### LC–MS/MS data processing

Waters.raw dataset were directly imported into Mzmine 2.30 (Pluskal *et al.*, 2010). The extracted ion chromatograms (XICs) were built with ions showing a minimum time span of 0.01 min, minimum height of 4000, and *m/z* tolerance of 0.001 (or 5.0 ppm). The chromatographic deconvolution was achieved by the baseline cut-off algorithm, with the following parameters: minimum peak height of 2500, peak duration range of 0.02–0.20 min, and baseline level of 500. Deconvoluted XICs were deisotoped using the isotopic peaks grouper algorithm with a *m/z* tolerance of 0.006 (or 10.0 ppm) and a retention time (t_R_) tolerance of 0.15 min. XICs were aligned together into a peak table, using the join aligner module (*m/z* tolerance at 0.006 (or 10.0 ppm), absolute t_R_ tolerance at 0.2 min, weight for *m/z* of 70, and weight for t_R_ of 30); ions from MS contaminants identified by blank injection and duplicate peaks were manually removed from the aligned peak table. The filtered peak table was eventually gap-filled with the peak finder module (intensity tolerance at 30.0 %, *m/z* tolerance at 0.001 Da (or 5.0 ppm), and absolute t_R_ tolerance of 0.2 min).

### LC–MS/MS data analyses

The preprocessed chromatograms were exported to GNPS (https://gnps.ucsd.edu) for molecular networking (Wang *et al.*, 2016). MS/MS spectra were window filtered by choosing only the top six peaks in the ± 50Da window throughout the spectrum. A network was then created where edges were filtered to have a cosine score above 0.70 and more than four matched peaks. Further edges between two nodes were kept in the network and only if each of the nodes appeared in each other’s respective top 10 most similar nodes. The spectra in the network were then searched against the spectral library of GNPS; the library spectra were filtered in the same manner as the input data. The molecular network was visualized using Cytoscape 3.5.1 (Shannon *et al.*, 2003). Peak area data from the Mzmine-processed LC–MS peaktable were combined with the spectral network, and visualized by plotting pie charts. Phylogenetic information was mapped on the network, by assigning unique colors to each genus. The constructed molecular network was further analyzed using the Network Annotation Propagation (NAP; accessible through the GNPS web-platform) tool for structural annotation of spectral nodes. NAP utilizes MetFrag i*n silico* fragmentation tool to search the structural databases of GNPS, Dictionary of Natural Products (DNP), and Super Natural II (Banerjee *et al.*, 2015). All precursor ions were hypothesized to be deprotonated molecular ions [M − H]^−^, and accuracy for exact mass candidate search was set to 10. *Fusion* and *Consensus* scores were calculated based on 10-first candidates in the network propagation phase.

The preprocessed LC–MS/MS peaklist file was also subjected to MS2LDA (https://MS2LDA.org) (Wandy *et al.*, 2017) for extracting MS2motifs. Parameters for the MS2LDA experiment were set as follows: input format MGF, *m/z* tolerance 5.0 ppm, t_R_ tolerance 10.0 s, minimum MS1 intensity 0 a.u., minimum MS2 intensity 50.0 a.u., no duplicate filtering, number of iterations 1000, number of Mass2Motifs 200.

All scripts used for data analyses platform integration are publically accessible at: https://github.com/DorresteinLaboratory/supplementary-Rhamnaceae.

### Data availability

LC–MS/MS raw data, the preprocessed peaklist file, and the integrated Cytoscape network file are deposited in the Mass spectrometry Interactive Virtual Environment (https://massive.ucsd.edu) with the accession number MSV000081805, which is accessible via the following link: https://massive.ucsd.edu/ProteoSAFe/dataset.jsp?task=36f154d1c3844d31b9732fbaa72e9284

The molecular network and NAP result of Rhamnaceae extracts can be found at the GNPS website with the following links:

https://gnps.ucsd.edu/ProteoSAFe/status.jsp?task=e9e02c0ba3db473a9b1ddd36da72859b
https://proteomics2.ucsd.edu/ProteoSAFe/status.jsp?task=6b515b235e0e4c76ba539524c8b4c6d8

The MS2LDA results are accessible through the following link: http://ms2lda.org/basicviz/summary/566; the summary for all Mass2Motifs from this study is available as Table S2.

## ACKNOWLEDGEMENTS

We thank the NIH for supporting this work under NIH-UCSD Center for Computational Mass Spectrometry P41 GM103484, and the NIH grants GMS10RR029121, R03 CA211211, and on tools for rapid and accurate structure elucidation of natural products R01 GM107550. This research was supported by the Basic Science Research Program through the National Research Foundation of Korea (NRF), which was funded by the Ministry of Science, ICT and Future Planning (NRF-2015M3A9A5030733) and also supported by Korea Research Institute of Bioscience and Biotechnology (KRIBB) initiative program of Republic of Korea. We thank the National Biodiversity Institute of Costa Rica (INBio), the National System of Conservation Areas and the Conservation Areas of Guanacaste, Arenal, and Central, Costa Rica, the Institute of Traditional Medicine, Laos, and International Biological Material Research Center (IBMRC) of KRIBB for providing the various plant extracts. KBK was supported by Sookmyung Women’s University Specialization Program Funding. JJJvdH gratefully acknowledges an ASDI eScience grant (ASDI.2017.030) from the Netherlands Organization for Scientific Research (NWO). MHM was supported by Veni Grant 863.15.002 from The Netherlands Organization for Scientific Research (NWO). We thank Prof. S. H. Kim (Yonsei University) for providing us standard compounds.

## AUTHOR CONTRIBUTIONS

KBK, ME, JJJvdH,MM, SHS, and PCD conceived the study. KBK and JP performed the LC–MS/MS analyses. KBK processed and analyzed the MS/MS data, and performed the manual inspection on experimental MS/MS data and the distributional analysis. RRdS developed the NAP annotation analysis. JJJvdH developed the MS2LDA topic modeling analysis. ME and JJJvdH developed the semi-automated annotation workflow by combining mass spectral molecular networking with MS2LDA, *in silico* annotation and ClassyFire. KBK, ME, and JJJvdH wrote the manuscript with discussion and help from all authors.

## SUPPORTING INFORMATION

Additional Supporting Information may be found in the online version of this article.

**Result S1.** Inspection on putative annotation of molecular families C and E.

**Result S2.** Validation of spectral identification using reference standards

**Figure S1.** NAP/MS2LDA-driven metabolite annotation of (a) flavonol 3-*O*-glycosides (molecular family **C**) and (b) cyclopeptide alkaloids (molecular family **E**).

**Figure S2.** Chromatographic validation for emodin-8-*O*-β-D-glucopyranoside (**9**).

**Figure S3.** Chromatographic validation for 3-*O*-protocatechuoylceanothic acid 2-methyl ester (**10**).

**Figure S4.** Chromatographic validation for 3-*O*-vanilloylceanothic acid (**11**).

**Figure S5.** Chromatographic validation for 3-*O*-protocatechuoylceanothic acid (**12**).

**Figure S6.** Chromatographic validation for nicotiflorin (**14**).

**Figure S7.** Chromatographic validation for quercetin 3-*O*-neohesperidoside (**15**).

**Figure S8.** Chromatographic validation for adouetine X (**18**).

**Figure S9.** Chromatographic validation for emodin (**21**).

**Figure S10.** ClassyFire consistency scores for the Rhamnaceae molecular network.

**Figure S11.** The complete chemical subclass distribution heatmap.

**Figure S12.** The complete Mass2Motif distribution heatmap.

**Table S1.** Detailed information about Rhamnaceae plant samples.

**Table S2.** The list of 200 Mass2Motifs extracted from Rhamnaceae dataset.

